# eTumorMetastasis, a network-based algorithm predicts clinical outcomes using whole-exome sequencing data of cancer patients

**DOI:** 10.1101/268680

**Authors:** Jean-Sébastien Milanese, Chabane Tibiche, Naif Zaman, Jinfeng Zou, Pengyong Han, Zhiganag Meng, Andre Nantel, Arnaud Droit, Edwin Wang

**Affiliations:** National Research Council Canada, 6100 Royalmount Ave, Montreal, Canada, H4P 2R2; Genomics Center, Centre Hospitalier Universitaire de Québec - Université Laval Research Center, Quebec, Canada, G1V 4G2; Department of Biochemistry & Molecular Biology, Medical Genetics, and Oncology, University of Calgary, 3330 Hospital Dr. NW, Calgary, Canada, T2N 4N1; Alberta Children’s Hospital Research Institute and Arnie Charbonneau Cancer Research Institute, University of Calgary, 3330 Hospital Dr. NW, Calgary, Canada, T2N 4N1; Chinese Academy of Agricultural Science, No. 12 Zhongguangcun South St., Haidian District, Beijing, 100086, China; Department of Medicine, McGill University, 3605 Mountain St, Montreal, Canada H3G 2M1

## Abstract

Continual reduction in sequencing cost is expanding the accessibility of genome sequencing data for routine clinical applications. However, the lack of methods to construct machine learning-based predictive models using these datasets has become a crucial bottleneck for the application of sequencing technology in clinics. Here we developed a new algorithm, eTumorMetastasis, which transforms tumor functional mutations into network-based profiles, and identify network operational gene signatures (NOG signatures) which model the tipping point at which a tumor cell shifts from a state that doesn’t favor recurrences to one that does. We showed that NOG signatures derived from genomic mutations of tumor founding clones (i.e., the ‘most recent common ancestor’ of the cells within a tumor) significantly distinguished recurred and non-recurred breast tumors. These results imply that somatic mutations of tumor founders are association with tumor recurrence and can be used to predict clinical outcomes. Finally, the concepts underlying the eTumorMetastasis pave the way for the application of genome sequencing in predictions for other complex genetic diseases.

## Introduction

As genome sequencing becomes cheaper and more convenient, it will become more accessible for routine clinical usage and its demand will rise. It has been expected that massive genome sequencing data combined with phenotypic and complex disease data will allow us to decode the underlying molecular mechanisms of diseases, and further to predict phenotypes and complex disease outcomes using genome sequencing data. Moreover, to fulfill the promises of precision medicine, it is necessary to construct clinically useful predictive models using DNA sequencing data. However, using these datasets to construct predictive models has become a crucial bottleneck in genomic biomarker development. Thus far, none of the existing machine learning algorithms is suitable to construct predictive models for diseases from genome sequencing data alone. For example, in a recent Dialogue for Reverse Engineering Assessment and Methods (DREAM) effort, scientists have tested more than 50 existing prediction algorithms and shown that none of them is able to construct cancer drug-responding predictive models using genome sequencing data alone^1^.

The huge challenge we are facing to construct predictive models using genome sequencing data is that complex diseases are often modulated by multiple distinct genetic pathways. For a given phenotype (*i.e.*, a complex disease), there are many ways to produce the phenotype, each of which is formed by the combined effects of multiple genes whose functions can be modulated through either genetic of epigenetic changes. Thus, different individuals who have the same phenotype/disease may have different causal genes and thus, may express different optimal drug targets. As an added level of complexity, tumors often exhibit extensive mutational heterogeneity, with genes’ mutation status varying widely across individual cancer cells. In another word, mutated genes are rarely shared between any two individual tumors of even a same cancer type and each patient has an individually unique genomic profile^2–7^. This feature of tumor mutations makes it extremely challenging to apply machine learning approaches for accurately predicting clinical outcomes based only on their genome sequences.

To cope with this challenge, we developed a novel network-based method to construct predictive models using the collective impact of genomic alterations in tumors, focusing on functionally mutated genes. We model the tipping point when a system shifts abruptly from one cellular state to another. As a proof-of-concept, we developed an algorithm, eTumorMetastasis, to construct predictive models of tumor recurrence using tumor whole-exome sequencing data, and showed that mutations in the tumor founding clones (i.e., the founding cancer cell transformed from a single normal cell through the acquisition of a series of mutations) can be used to predict tumor recurrence in breast cancers. Finally, we envisioned that this method could be widely applicable to construct predictive models for other complex genetic diseases and phenotypes using genome/whole-exome sequencing data.

## Results

### An overview of the eTumorMetastasis

Tumor recurrence and metastasis is the leading cause of cancer mortality and an accurate evaluation of this process could greatly aid clinicians in making treatment decisions. For example, most of the low-risk breast cancer patients (i.e., patients whose tumors would not recur for 10 years after surgery alone) do not gain survival benefits from adjuvant therapy (i.e., chemotherapy given after surgery to reduce the risk of cancer recurrence), but will suffer from its toxic side effects. Prognostic biomarkers could predict whether a patient is more likely to suffer from tumor recurrence and metastasis and whether they would benefit from adjuvant chemotherapy.

**Fig 1.**
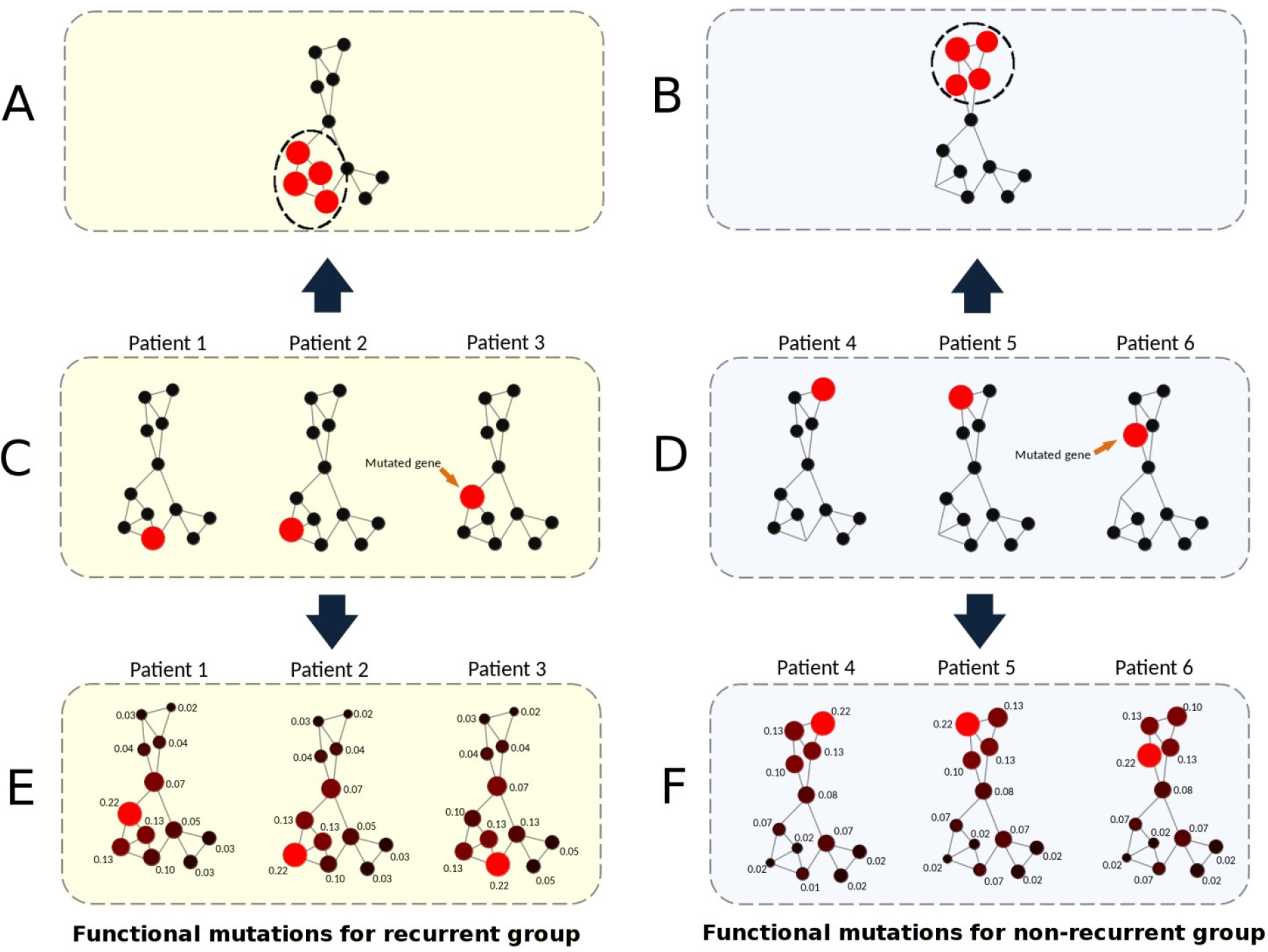
Network propagation and netProfiles for recurred and non-recurred samples. Functionally mutated genes in three recurred samples (**C**) are different but for a network cluster (**A**) in the human signaling network, and so are they in three non-recurred samples (**B, D**). For each recurred sample, by conducting network propagation based on its mutated genes, a network cluster (nodes in red and cyan, **E**) which is similar to the cluster in A is emerged. A similar pattern (**F**) is overserved for the non-recurred samples. The network clusters (**E,** they are similar between the recurred samples) and (**F,** they are similar between the non-recurred samples) make it possible to classify recurred and non-recurred samples, respectively. In the network, nodes and lines represent genes and gene relations, respectively. Numeral numbers of each network node represent heating scores. Red nodes represent mutated genes while cyan nodes represent gaining ‘energy’ and node sizes represent ‘hearting scores’ values - the amount of the gained energy.

To develop predictive models for cancer prognosis, machine learning requires the identification of features that distinguish recurred and non-recurred cancer patient groups: i.e., genes which are frequently functionally inactivated/activated within the recurred group but not in the non-recurred group, and vice visa. Because the mutation profiles of tumors at the gene level are sparse (Fig 1C, 1D), this type of data by itself is not suitable for machine learning approaches. However, we and others previously showed that functional cancer mutations collectively affect several network regions or subnetworks of the human signaling network (Fig 1A, 1B)^4,20,21^. These results suggest that cancer signaling processes triggered by the mutations of the recurred (or non-recurred) samples are convergent to several subnetworks or network regions (Fig 1E, 1F). Consequently, while recurred (or non-recurred) samples share certain impaired signaling processes/subnetworks, they do not necessarily share the same sets of mutated genes (i.e. there are many ways to “break” a subnetwork). These network regions or clusters may represent key cancer signaling processes underlying the molecular mechanisms of the samples in either the recurred or non-recurred group. Therefore, we envisioned that if we could use a tumor’s mutations to infer which network regions or clusters are functionally impaired, then we can identify shared features within the tumor samples of either the recurred or the non-recurred group.

Computational techniques such as random walk and network propagation enable us to transform mutations of a tumor into the perturbed signaling network regions or clusters in the human signaling network. The network propagation algorithm works by projecting the mutated genes of a tumor onto a cancer type-specific metastasis network in which each mutated gene is represented as a heat source. The heat source diffuses to neighboring genes along the edges of the network in a process which is analogous to heat diffusion. After a certain time period, the diffusion stabilizes to a point where each gene in the network will have received a certain amount of ‘energy’, which is represented by a ‘heating’ score (i.e., we called this process as network profiling here, while the resulting heating scores for all the network genes for a tumor sample is called netProfile). The value of the heating score of a gene could be treated in a similar fashion as a transcript abundance value for that gene. The higher a heating score is, the more functionally active (or inactive) the gene is. Thus, we expect that the genes in the common network regions of either the recurred or non-recurred group will have higher but similar heating scores (Fig 1D, 1E). Further, we appended the netProfile of each tumor to form a matrix (i.e., netMatrix) where network genes are rows and tumor samples are columns, and the values are heating scores. By doing so, we transformed a sparse gene mutation dataset into a data-richer matrix containing data similar to gene expression profiles.

From a systems biology perspective, a signaling network has a critical transition threshold (i.e., a tipping point) at which point the system shifts abruptly from one state to another ^26^. The critical transition threshold for a metastasis network could be marked by an abrupt change between the cellular states that favor or do not favor tumor recurrence^25^. Although cancer driver-mutated genes are highly diverse between tumors, for a recurred tumor, their collective effects on the metastasis network could converge to trigger a state’s switch (eg, leading to a phenotypic switch) from the non-recurring state to the recurrence-promoting state. We thus propose a ‘network operational gene signature’ (i.e., NOG signature) to quantify the two cellular states and the state’s switch. A NOG signature contains a set of genes whose heating scores (i.e., the mean of the scores) in the recurred samples and non-recurred samples represent the recurring and non-recurring cellular states, respectively. Further, these scores are not only far from the tipping point (i.e., state switch, we assume that the switch is the middle point of the distance between the two states) between the recurred and non-recurred states, but also significantly different between the two groups.

To identify the NOG signatures, we could apply the netMatrix algorithms that have historically been used for classifications based on gene expression profiles. However, unlike gene expression profiles, gene heating scores in the netMatrix are too weak to produce high-quality NOG signatures from traditional machine learning algorithms. Therefore, we modified our previously developed Multiple Survival Screening (MSS) algorithm^2^ to successfully identify NOG signatures which significantly distinguished recurred and non-recurred tumors. Finally, for a given new tumor sample’s whole-exome sequencing data, we calculate a netProfile and correlate it with the heating score profiles of the two states of the NOG signatures to successfully predict its prognosis. For example, if the netProfile is far from the tipping point and close to the heating score profile of the recurred state for a NOG signature, then we would assign that tumor to the recurred group. The implementation of these ideas (i.e., eTumorMetastasis) will be described in the next section.

### Tumor founding clone mutations predict tumor recurrence

To test if genome/whole-exome sequencing data can be used to robustly predict cancer prognosis, we used mutations identified in the tumor founding clones. A single normal cell could acquire a set of random mutations that allows it to be transformed into the founding cancer cell (e.g., the founding clone), an early evolutionary stage of a tumor. Additional accumulation of mutations from this founding clone leads to the formation of a tumor that is composed of a heterogeneous mixture of cells (Fig 2). Therefore, each tumor originates from a founding clone whose mutations are ubiquitously present in all the cells of that tumor. New mutations do not arise in isolation but rather act together in a complementary manner with the established genomic landscape; therefore, the pre-existing mutations or genetic variants of a molecular network may have a profound effect on cellular fate and determine whether novel mutations will result in altered cell death modulation, clonal expansion, metastasis and other cancer hallmark traits. This implies that the genetic makeup of the founding clone provides an evolutionary constraint for sequential subclones and limits the genetic and clonal complexity of tumors. Tumor genome sequencing and the frequencies of the observed mutations allow us to dissect founding and subclones’ mutations and to replay the tape of the tumor’s evolutionary history, while eTumorMetastasis (see Methods in Supplementary Materials) allows us to decipher the tumor evolution and patient outcomes that are affected by these early mutational events. Therefore, we examined whether the sum of somatic mutations in the founding clones could predict tumor recurrence and metastasis.

**Fig 2.**
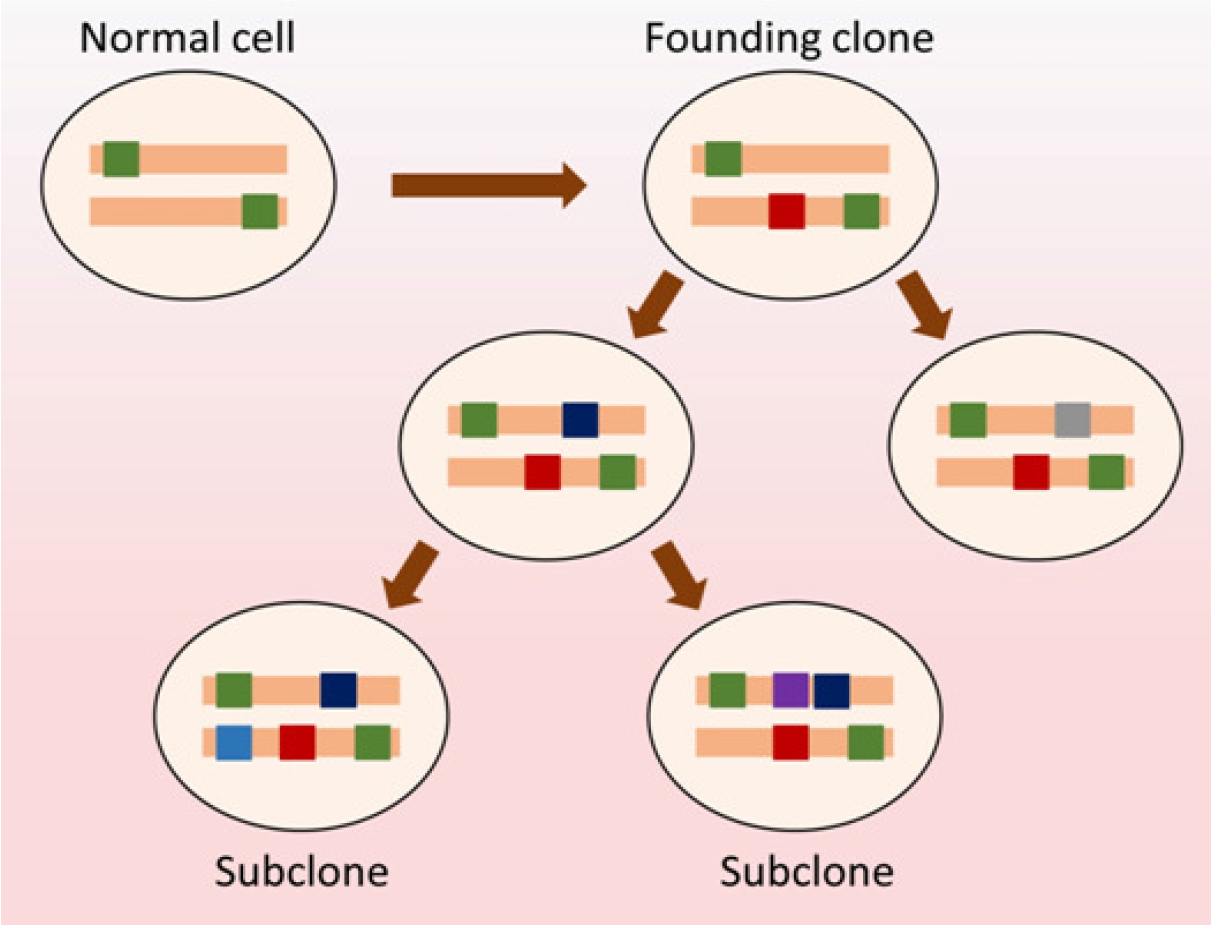
Tumor evolution, a founding clone and its mutations. New somatic mutations (dark brown) functionally or epistatically work with germline mutations (green) to form a founding clone (i.e., the earliest, ancestral cancer cell). New somatic mutations occur in the founding clone to generate subclones. A tumor often contains several subclones. Of note, all the mutations from germline and somatic mutations in the founding clone are present in all the subclones and every cancer cell of the tumor. Circles represent cells while colored bars represent mutated genes.

Fig 3 shows the flowchart of the eTumorMetastasis. Briefly, somatic mutations were identified using the whole-exome sequencing data of tumors and their paired normal samples. Tools such as CADD were applied to the mutations to identify functionally mutated genes (Fig 3A). Meanwhile, a cancer-specific metastasis network was constructed using gene expression data associated with cancer recurrence and a literature-curated signaling network (Fig 3B). The functionally mutated genes were seeded onto the network to initiate a network propagation that generates ‘heating scores’ for the network genes (Fig 3C). The ‘heating scores’ for the network genes from all the samples were then aggregated into a netMatrix to identify NOG signatures (Fig 3D, 3E). The details of the eTumorMetastasis implementation are described in the Supplementary Materials.

To examine whether the sum of somatic mutations in founding clones could predict tumor recurrence and metastasis, we used breast tumor whole-exome sequencing data. Breast cancer has two major subtypes: ER+/luminal and ER−/basal with ER+ tumors representing the largest proportion (~70%) of breast tumors. Through an examination of the sequencing data and clinical follow-up information, we found that TCGA collected several hundreds of ER+ tumors but only ~100 ER−/basal tumors. We therefore decided to use only ER+ tumors in this study. For each sample, we identified functionally mutated genes in the founding clones using the sequencing data of the breast tumors and their paired normal samples (Supplementary Materials). We then applied eTumorMetastasis (see Methods in Supplementary Materials) as shown in Fig 2 to identify NOG signatures that are necessary to predict tumor recurrence through a modification of the MSS algorithm^2^ (see Supplementary Materials).

**Figure 3.**
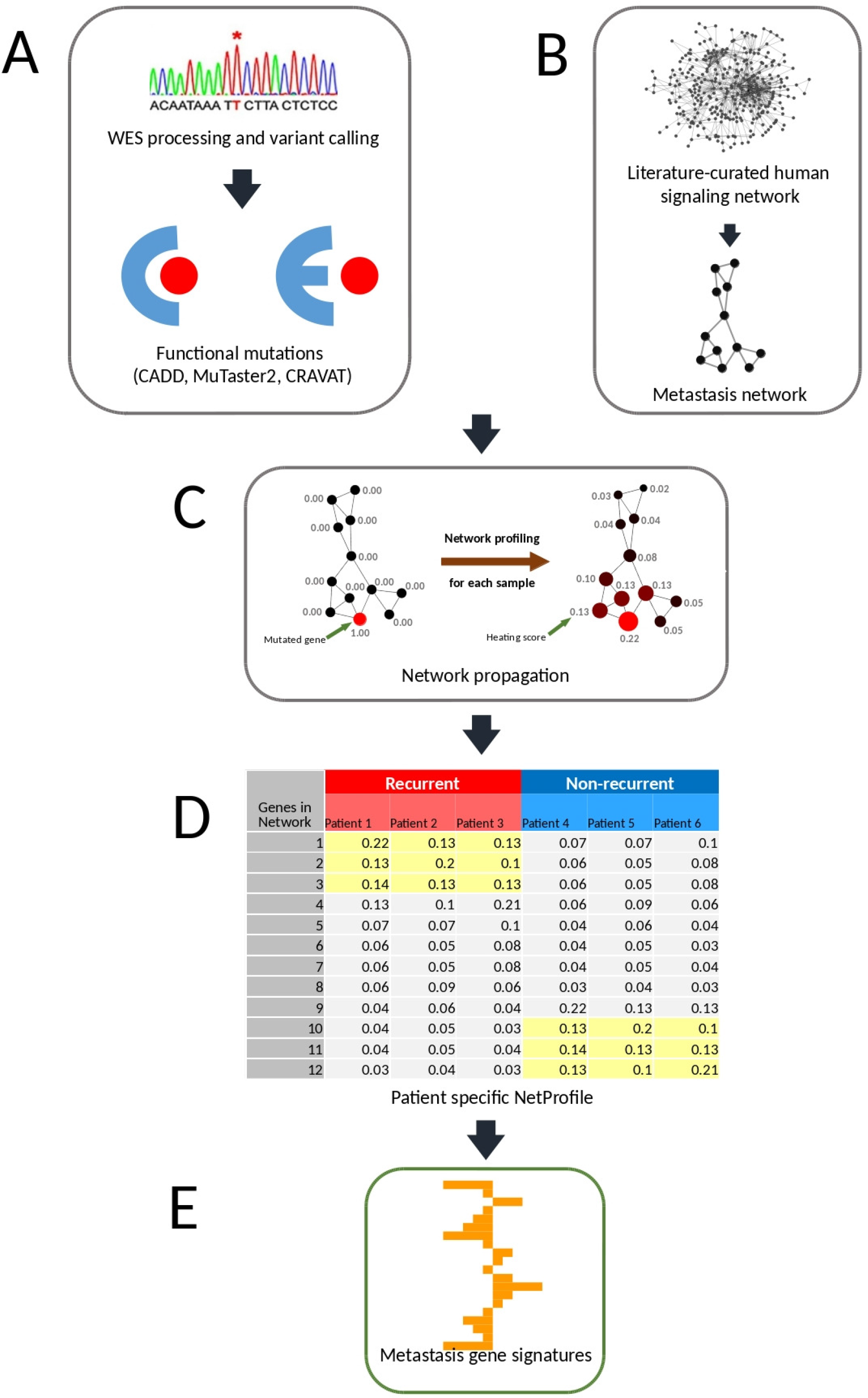
A flowchart of the eTumorMetastasis. (A) Somatic mutations are identified using whole-exome sequencing data of tumors and their paired normal samples, (B) A cancer-specific metastasis network is constructed using the gene expression data associated cancer recurrence and the literature-curated signaling network, (C) Functionally mutated genes of samples (eg, from tumor founding clones) are projected on a metastasis network to generate gene heating scores using a network propagation approach (i.e., mutated genes become heat sources that diffuse ‘energy’ to neighboring genes along the edges of the network), (D) Following network propagation, the ‘heating scores’ for the network genes for all the samples are aggregated into a matrix, called netProfile, and (E) The modified Multiple Survival Screening (MSS) algorithm is applied to the netProfile to identify gene signatures that distinguish the two phenotypic groups (i.e., recurred vs non-recurred tumor samples). Each signature containing a set of genes with heating scores quantifies the network state transition (i.e., state between recurred and non-recurred tumors). In the network, nodes and lines represent genes and gene relations, respectively. Numeral numbers of each network node represent heating scores. Red nodes represent mutated genes while cyan nodes represent gaining ‘energy’ and node sizes represent ‘heating scores’ - the amount of the gained energy.

We identified 18 NOG signatures and showed that these were significantly predictive in 2 validation sets (Supplementary Table 4). To further improve prediction accuracy, we built combinatory gene signature sets (NOG_CSS sets, Supplementary Methods and Supplementary Table 5)^3^ using the identified NOG signatures. We showed that founding clone-derived NOG signatures significantly distinguished recurred and non-recurred tumors in the validation sets of ER+-breast cancer patients (Table 1, Fig 4A). We further showed that the NOG_CSS sets (Table 1) significantly distinguished recurred and non-recurred tumors in an additional validation set (P=7.94×10^−4^, Table 1 and Fig 4B, 200 samples). Finally, we validated these NOG gene signatures using the independent TCGA-CPTAC set which contains 295 additional ER+ breast cancer samples (P=0, Table 1, Fig 4C).

**Table 1.** Prediction accuracy and recall rate for validation sets for breast cancer using the NOG_CSS sets derived from tumor founding clones.

| Dataset | Number of samples | Low-Risk |  | High-risk |  |
| --- | --- | --- | --- | --- | --- |
|  |  | Accuracy (%)<sup>*</sup> | Recall (%)<sup>†</sup> | Accuracy (%)<sup>**</sup> | Recall (%)<sup>††</sup> |
| TCGA-Nature | 200 | 94.3 | 46.1 | 30.4 | 35.0 |
| TCGA-CPTAC | 295 | 95.1 | 73.6 | 52.0 | 38.2 |

Notes:
\*Percentage of non-recurred (i.e., non-metastatic) samples in the predicted low-risk group.
†Percentage of the predicted low-risk samples from the non-recurred group.
\*\*Percentage of recurred (i.e., metastatic) samples in the predicted high-risk group.
††Percentage of the predicted high-risk samples from the recurred group.

**Fig 4.**
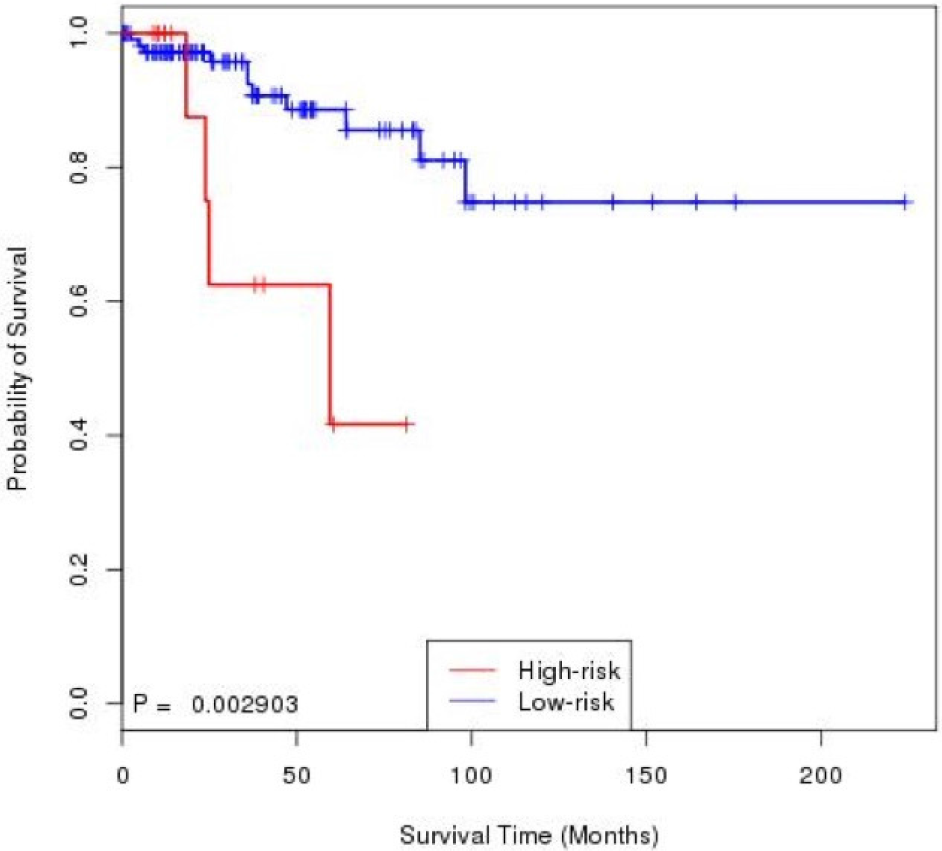
Kaplan–Meier curves of the risk groups for breast cancer patients with 10-year disease-free survival predicted by the NOG_CSS sets. NOG_CSS sets derived from tumor founding clones’ mutations (i.e., somatic and germline mutations) in (**A**) the training set, (**B**) the validation set, TCGA-Nature and (**C**) the validation set, TCGA-CPTAC.

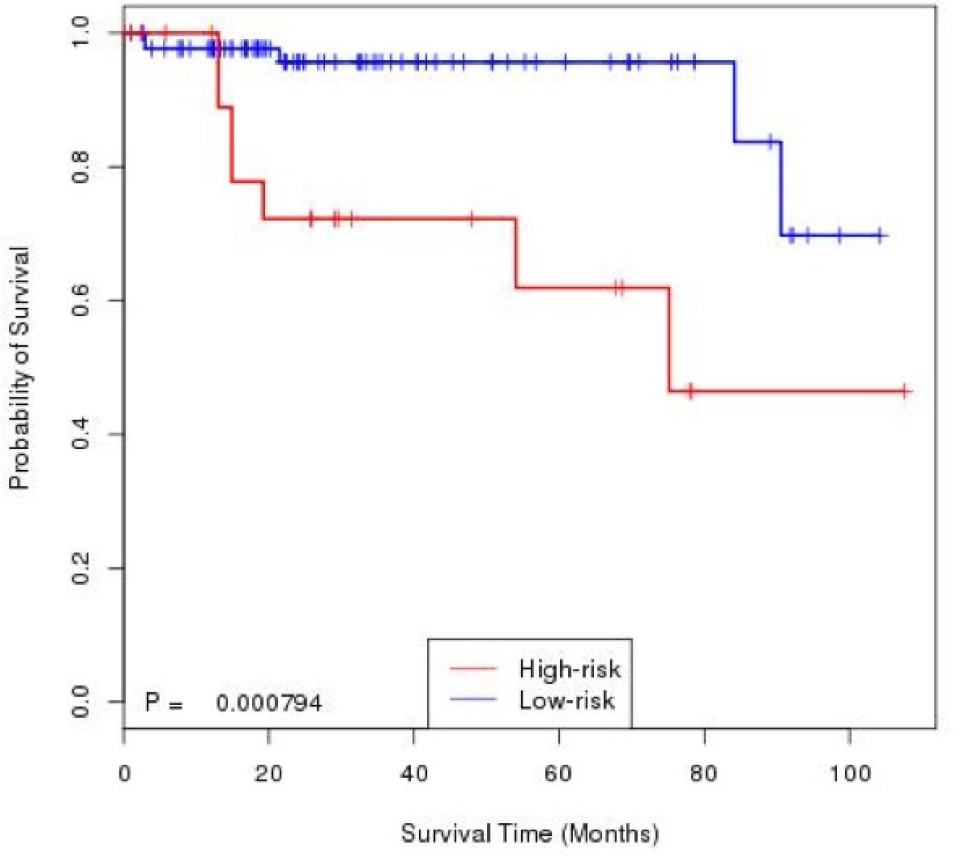

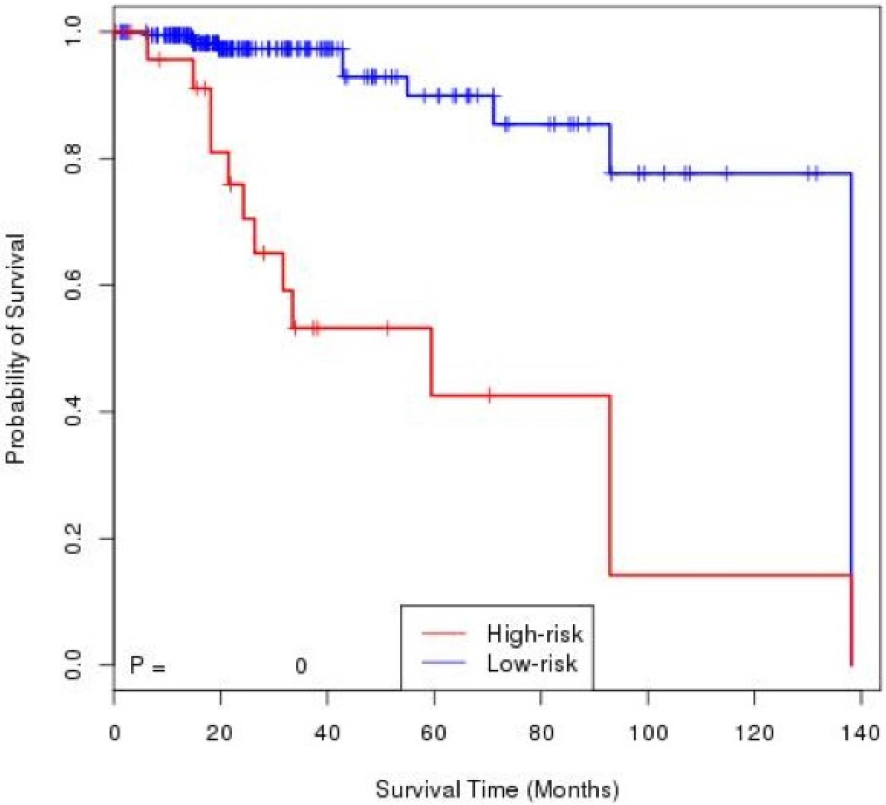

## Discussion

### Predicting cancer prognosis using genome sequencing data

Genome sequencing technologies are already being used for personalized genomic tests such as TruGenome clinical sequencing tests (Illumina), Personal DNA tests (23andMe), FoundationOne cancer sequencing tests (Foundation Medicine) and so on. At the moment, it is easy to generate personal genome sequencing data, but extremely difficult to interpret them in the context of personal health. Genomic reports derived from these genomic tests often use single gene mutation or SNP to estimate risks of complex genetic diseases or to guide certain treatments. However, most of these associations are very weak and ambiguous because complex genetic diseases are derived from the interactions of multiple genetic mutations and environmental factors. Clearly, to properly interpret personalized genomic test results, it is preferable to use a collection of mutations to predict phenotypes or diseases. Tumor genome sequencing provides a rich new source of data for constructing predictive models but has proven difficult to use for that purpose because tumors rarely share the same driver-mutating genes, even those the same cancer type ^2,26^. In this study, we showed that it is possible to predict cancer prognosis using genome sequencing data from multiple genes, which paves the way for improved interpretation of sequencing technology on clinical samples.

We demonstrated that the examination of the collective effects of genomic alterations (i.e., groups of functionally mutated genes) on cancer hallmark networks are more capable of representing the phenotypic consequences of mutated genes. Rare commonalities of mutated genes between tumors suggest that there are multiple ways in which genetic alterations may trigger such a state transition. However, the collective effect of multiple mutated genes may converge into a few sets of networked genes (i.e., NOG gene signatures) that modulate the transition state between metastasis and non-metastasis. Each of these sets (i.e., NOG gene signatures) contains a group of genes encoding their regulatory relationships and strengths (i.e., represented by heating scores here). In this context, we expected that these NOG gene signatures might predict tumor recurrence and metastasis.

Because cancer driver-mutated genes are sparse among tumors, the identification of the NOG gene signatures requires us to modify this data to estimate their collective impacts on signaling and functional networks. Therefore, we applied a network profiling approach to diffuse the effects of these functionally mutated genes on networks so that we could identify the subnetwork that are commonly impacted in either the recurred or the non-recurred groups (Fig 1). These common network regions provide the means for extracting common features (i.e., genes with similar heating scores) in one group but not in the other group. Of note, in the past, the network propagation algorithm has been mainly used for network topological modeling^24^. It has been used in the HotNet algorithm^16^ to identify significantly mutated subnetworks in the TCGA data and classify tumor subtypes^11,25^. Finally, we showed that NOG_CSS sets derived from the NOG gene signatures significantly improved the performance of predictive models constructed from genome sequencing data. The concepts here could be widely applicable to other complex diseases for constructing predictive models using genome sequencing data.

In summary, it is possible to predict cancer prognosis based on genome sequencing data, which paves the way to the application of genome sequencing technology in the clinics. We showed that sequencing of patient founding clones might provide an efficient and convenient way for predicting tumor recurrence. Finally, the concepts for developing the eTumorMetastasis could be used for predicting clinic outcomes of other complex genetic diseases using genome sequencing data.

## Methods

### Data for tumors and paired normal samples

Whole exome-sequence data of ER+ breast tumors and their paired normal samples were obtained from TCGA (The Cancer Genome Atlas). Patients who have information on recurrence and clinical follow-up were selected for the analysis. Based on these criteria, we retained 400 ER+ breast tumor samples (released by TCGA in 2012, we called it TCGA-Nature set). Further, we obtained data from 295 ER+ breast tumor samples in the cBioPortal (http://cbioportal.org) which was released in 2017 by TCGA-CPTAC (The Clinical Proteomic Tumor Analysis Consortium). We called these the TCGA-CPTAC set. In total, data from 695 (i.e., a TCGA set, a TCGA-CPTAC set and randomly selected 200 samples from the TCGA set for training) ER+ breast tumor samples were used in this study (Supplementary Tables 1 and 2). Gene expression profiles and SNP 6.0 array data from these samples were also downloaded from TCGA and cBioPortal. The samples/data were processed following examination of the tumor purity and variant calling (Supplementary Methods). The sample numbers remaining after each processing step in the training and validation sets have been listed in Supplementary Table 2.

### Determination of functionally mutated genes

To determine whether a genetic variant is functionally mutated, we applied the following tools: CRAVAT^17^ (e.g., a functional mutation is defined as a score higher than or equal to 0.5), MutationTaster2^18^ (e.g., a functional mutation is defined as having a disease impact) and CADD (e.g., Combined Annotation Dependent Depletion, used C-score 10 or CADD-10 as a cutoff which is suggested by the authors). For a given sample, we merged all functional mutations predicted by the three tools for further analysis. In this study, all the mutated genes mentioned are ‘functionally mutated genes’. In each founding clone, the average number of the somatic mutation in coding regions and the functionally mutated genes defined by each tool is listed in https://github.com/WangEdwinLab/eTumorMetastasis.

### Construction of ER+ breast cancer-specific metastasis networks

To construct a breast ER+ breast cancer specific metastasis network, we modified the procedure for constructing breast ER+ specific cancer survival and proliferation networks^20^. Briefly, we extracted a subnetwork by mapping the ER+ breast cancer specific metastasis-associated genes onto the literature-curated human signaling network. To do so, we first identified metastasis-associated genes using ER+ cancer cell lines and tumor samples. We obtained the gene expression data of 22 ER+ cancer cell lines from the Cancer Cell Line Encyclopedia (CCLE, http://www.broadinstitute.org/ccle). Gene expression data normalization was conducted with the median centering and z-score normalization method described previously^2^. Using the ratio of two genes’ expression values (i.e., CDH1/VIM for determining epithelial–mesenchymal transition), we classified these lines into epithelial (n=13, CDH1/VIM >1.2) and mesenchymal (n=9, CDH1/VIM <= 1.2) lines. Modulated genes (called Set 1, see https://github.com/WangEdwinLab/eTumorMetastasis) were identified by conducting t-test (P < 0.05) comparison of the two groups of cell lines with 10 re-samplings (i.e., each re-sampling randomly took 60% of the original samples). We also obtained the gene expression data of 1,197 ER+ tumor samples from METBRIC set (https://www.ebi.ac.uk/ega/studies/EGAS00000000083). These samples have information about cancer recurrence and clinical follow-up. Using this set, we used a t-test (P < 0.05) to identify the modulated genes between the recurred and non-recurred samples. Next, we performed Kaplan-Meier survival tests on the modulated genes to identify survival-associated genes (called Set 2, see https://github.com/WangEdwinLab/eTumorMetastasis) using 10 resamplings. We further identified potential cancer gene regulators by analyzing the copy number data (SNP 6.0 data) of the ER+ tumor samples from TCGA. The SNP 6.0 data were processed using GISTIC to obtain GISTIC scores for each gene. For a given gene in a sample, if its GISTIC score was greater than 0.3 and its expression value was ranked among the top 50% of the genome, we defined this gene as a cancer regulator (for details, see Zaman et al., 2013^20^). The regulators of all TCGA’s ER+ tumor samples were defined as Set 3 (see https://github.com/WangEdwinLab/eTumorMetastasis). We mapped all the genes from Sets 1, 2 and 3 onto the signaling network (eg, merging the manually curated human signaling network and a protein interaction network and extracted their links (i.e., we kept the network genes which are common with the genes in Sets 1, 2, or 3, and their links in the network and then removed the other genes and their links) to obtain an ER+ breast cancer specific metastasis network that contains 6,148 genes and 62,004 interactions (see https://github.com/WangEdwinLab/eTumorMetastasis).

### Generating netProfiles using a network propagation approach

To generate a netProfile for a sample, we projected its mutated genes as seeds onto a cancer type-specific metastasis network and then applied the network propagation algorithm^24^ to obtain heating scores of the genes within the network. We applied a scaling factor of 100,000 to the heating scores and then conducted data transformation using the median centering and z-score within sample approach^2^. To examine the potential effects of the scaling factor, we conducted a sensitivity analysis by re-running the eTumorMetastasis using the scaling factor of 10,000 after the network propagation. We found that the prediction accuracies of the gene signatures are similar to those of the gene signatures which are obtained by using the scaling factor of 100,000.

### Statistical Analysis

Statistical significance of the prognostic groups (ie, high- or low-risk groups defined by gene signatures) was determined using Kaplan-Meier survival plots. A prognostically significant result was defined by log-rank P< .05. Prognostic significance of clinicopathologic factors and molecular features (i.e., mutated genes) were performed with the use of the Cox proportional hazards regression model. P values were based on likelihood ratio tests. All the analyses were performed using the statistical R package.

